# Long dsRNA mediated RNA interference (dsRNAi) is antiviral in interferon competent mammalian cells

**DOI:** 10.1101/2022.01.14.476298

**Authors:** SL Semple, RA Jacob, K Mossman, SJ DeWitte-Orr

## Abstract

In invertebrate cells, RNA interference (RNAi) acts as a powerful defense against virus infection by cleaving virally produced long dsRNA into siRNA by Dicer and loaded into RISC which can then destroy/disrupt complementary viral mRNA sequences. Comparatively in mammalian cells, the type I interferon (IFN) pathway is the cornerstone of the innate antiviral response. Although the cellular machinery for RNAi functions in mammalian cells, its role in the antiviral response remains controversial. Here we show that IFN competent mammalian cells engage in dsRNA-mediated RNAi. We found that pre-soaking mammalian cells with concentrations of sequence-specific dsRNA too low to induce IFN production could significantly inhibit viral replication, including SARS-CoV-2. This phenomenon was dependent on dsRNA length, was comparable in effect to transfected siRNAs, and could knockdown multiple sequences at once. Additionally, Dicer-knockout cell lines were incapable of this inhibition, confirming use of RNAi. This represents the first evidence that soaking with gene-specific dsRNA can generate viral knockdown in mammalian cells. Furthermore, demonstrating RNAi below the threshold of IFN induction has uses as a novel therapeutic platform.

## 1. Introduction

In their historic discovery of the DNA double-helix, Watson and Crick openly stated that a similar molecule derived from ribose was “probably impossible” due to its anticipated stereochemistry (Watson & Crick, 1953). Remarkably, just three years later, this sceptical view of double-stranded RNA (dsRNA) was disproven by Rich & Davies (1956) when polymers of polyadenylic acid were shown to hybridize with polyuridylic acid to produce diffraction patterns typical of a helical structure. This breakthrough would revolutionize techniques in molecular biology, but this was not the only field that would be heavily influenced. Because it was known that some viruses contained only RNA, the newfound double-stranded potential provided an answer for how this nucleic acid, and hence viruses, could replicate. Scientists now realize that essentially all viruses produce long dsRNA (>40 bp) at some point during replication and that this molecule is not found in normal, healthy cells (Weber *et al*., 2006; Son *et al*., 2015). Long dsRNA acts as a pathogen associated molecular pattern (PAMP) and is capable of alerting host immune defenses to viral infection (reviewed by DeWitte-Orr and Mossman, 2010). When dsRNA binds to its complementary pattern recognition receptors (PRRs), several downstream responses activate host antiviral immunity (reviewed by Jensen and Thomsen, 2012). Interestingly, the resultant antiviral immune response can vary significantly depending on whether the host is a vertebrate or not.

When long extracellular dsRNA is detected in a vertebrate host, rapid induction of the type I interferon (IFN) pathway occurs. In this scenario, extracellular dsRNA first binds to class A scavenger receptors (SR-As) located on the cell membrane (DeWitte-Orr *et al*., 2010). The dsRNA is then taken up through receptor mediated endocytosis and remains in the endosome until it either binds to an endosomal PRR, called toll-like receptor 3 (TLR3, reviewed by Matsumoto *et al*., 2014), or is transported into the cytoplasm via the SIDT2 molecular channel (Nguyen *et al*., 2017). Once in the cytoplasm, the dsRNA is free to interact with cytoplasmic PRRs known as retinoic acid-inducible gene-I (RIG-I)-like receptors (RLRs) (reviewed by Rehwinkel and Gack, 2020). Regardless of their location, successful binding of dsRNA to PRRs induces a signaling cascade resulting in the production of type I IFNs, primarily IFNαs and IFNβ (reviewed by Li *et al*., 2018). These antiviral cytokines can then act in an autocrine or paracrine manner by binding to their cognate receptors which induces expression of IFN-stimulated genes (ISGs), such as CXCL10 (reviewed by Borden *et al*., 2007; Cheon *et al*., 2014). Proteins encoded by ISGs can function to limit viral infection both directly, through inhibition of translation, and indirectly, by enhancing the adaptive immune response towards viral pathogens (reviewed by Yang and Li, 2020). This broad-spectrum “antiviral state” results in a slowed metabolism and the eventual apoptosis of infected cells, hence limiting viral spread (reviewed by Fritsch and Weichhart, 2016). Importantly, the potency of the type I IFN pathway is dependent on the length of dsRNA molecules, but sequence does not appear to influence this response (Kato *et al*., 2008; Leonard *et al*., 2008; DeWitte-Orr *et al*., 2009; Poynter and DeWitte-Orr, 2018). As a result, the IFN pathway is recognized to induce powerful, yet non-specific, inhibition of viral replication.

In contrast, RNA interference (RNAi) is the main antiviral mechanism used when invertebrate cells encounter viral dsRNA. In the early stages of this response, RNAi appears very similar to the IFN pathway. Long extracellular dsRNA is brought into the cell either by SR-As into endosomes or transported directly into the cytoplasm via the molecular channel SID-1 (Ulvila *et al*., 2006; Winston *et al*., 2002). Similar to its mammalian homolog SIDT2, SID-1 in invertebrates transports dsRNA in a length-dependent and sequence-independent manner (Li *et al*., 2015). Once within the cytosol, the dsRNA is cleaved into small interfering RNAs (siRNAs) by Dicer, a dsRNA-specific RNase-III-type endonuclease (reviewed by Maillard *et al*., 2019). A single strand of each siRNA duplex is then bound by Argonaute (Ago), which combines with accessory proteins to form the RNA-induced silencing complex (RISC, reviewed by Maillard *et al*., 2019). The siRNA-loaded RISC acts to render complementary target RNAs useless, either by mediating their cleavage or by remaining bound to prevent their translation (reviewed by van den Berg *et al*., 2008). Contrary to the IFN pathway, the RNAi pathway is heavily dependent on complementarity of sequence between the siRNA and the cytosolic target RNA.

Though mammalian cells have been shown to possess all the cellular machinery needed for RNAi, it is currently believed that these organisms only use the IFN pathway to combat viral invaders (reviewed by Schuster *et al*., 2019). Mammalian cells can undergo RNAi with long dsRNA (dsRNAi) when IFN-incompetent or when IFN competent and transfected with siRNA (Elbashir *et al*., 2001; Billy *et al*., 2001; Yang *et al*., 2001; Paddison *et al*., 2002; Maillard *et al*., 2013; Maillard *et al*., 2016). When mammalian cells have normal IFN function and are exposed to long dsRNA, it has been shown that the IFN pathway actively inhibits RNAi (Seo *et al*., 2013; Van der Veen *et al*., 2018). However, none of these studies soaked cells with long dsRNA at concentrations that were too low to induce the IFN response. Moreover, many of these studies either transfect cells with long dsRNA or use the TLR3 agonist, polyinosinic:polycytidylic acid (pIC, reviewed by Komal *et al*., 2021). The use of pIC is an excellent tool for understanding the IFN pathway, but it is important to note that this molecule is not the same as naturally occurring dsRNA. It has no defined length, a preparation of pIC can range from 1.5 kb to 8 kb and contains complimentary strands of inosines and cytosines, that would be not found in nature, to produce a dsRNA helix (Scadden, 2007). Thus, the results from both pIC and dsRNA transfection studies may not be indicative of the natural cellular responses to extracellular dsRNA, particularly at low concentrations. This suggests a fascinating possibility, where RNAi is the mechanism used by mammalian cells when dsRNA levels are too low to induce IFN. This *sentinel* activity could provide pre-emptive protection and/or clearance early in the course of infection when viral numbers are not yet high enough to warrant the costly use of the IFN pathway.

Since its discovery in 1998 by Fire and colleagues, scientists have been fascinated with the gene knockdown potential of RNAi. Yet, as described above, this sequence-specific knockdown did not seem possible in IFN-competent mammalian cells without the use of transfection agents. Moreover, the understanding of how cells respond to non-IFN inducing concentrations of dsRNA is completely absent from the literature. In the present study, we provide evidence that antiviral RNAi can be induced in mammalian cells by simply pre-soaking the cells with dsRNA at concentrations that are too low to induce IFN production. Remarkably, we were able to demonstrate this phenomenon in multiple mammalian cell types using several different dsRNA sequences to inhibit the infection of vesicular stomatitis virus expressing green fluorescent protein (VSV-GFP), as well as the human coronaviruses (CoV) HCoV-229E and SARS-CoV-2. Additionally, we reveal that this phenomenon is length-dependent and requires the presence of Dicer. Aside from the implications this work could have on developing novel antiviral/gene therapies, these results provide an explanation as to why the mammalian lineage retained all the necessary machinery for RNAi and why several mammalian viruses have devoted parts of their valuable genetic material to inhibit this pathway (Wang *et al*., 2006; Yang *et al*., 2013; Qui *et al.*, 2020).

## 2. Results

### 3.1 The interferon response in cells soaked with long dsRNA versus pIC

Because RNAi appears to be masked by the interferon response, it was crucial to identify which concentration of soaked dsRNA would not induce the IFN pathway. When gene expression of IFNβ and CXCL10 was measured 26h after cells were exposed to 700 bp GFP dsRNA (0.5 μg/mL and 10 μg/mL) or 10 μg/mL of HMW pIC, only the pIC condition appeared to induce the IFN response (**Figure 1**). The THF and SNB75 cell lines were initially selected for this study to explore whether both a normalized cell line (THF) and an “abnormal” cancerous cell line (SNB75) would be capable of long dsRNAi while being IFN competent. In THF, the gene expression of IFNβ increased only in the pIC exposure condition, but due to variability this was not significantly different from the other conditions (**Figure 1Ai**). When SNB75 was stimulated with these dsRNA and pIC doses, only the pIC treatment was able to induce significant upregulation of IFNβ gene expression (**Figure 1Bi**). Because IFNβ gene expression is known to be quite rapid and short-lived, the more persistent ISG, CXCL10, was also measured. In both THF and SNB75, CXCL10 gene expression was only observed to significantly increase when cells were soaked with pIC (**Figures 1Aii** and **1Bii**). Neither dsRNA concentration appeared to induce significant upregulation of IFNβ or CXCL10 when compared to the unstimulated control (**Figure 1**). When comparing the molar amounts, 5.1 nM of pIC (average length of 3000 bp) and 21.6 nM of 700 bp dsRNA was added to the cells. This means that four times more dsRNA molecules were added to each cell when compared to the number of pIC molecules._Furthermore, pIC is over four times longer than the dsRNA molecules used here, so it is difficult to compare efficacy of IFN induction between these molecules. As such, pIC should only be considered a positive control in this experiment.

**Figure 1:**
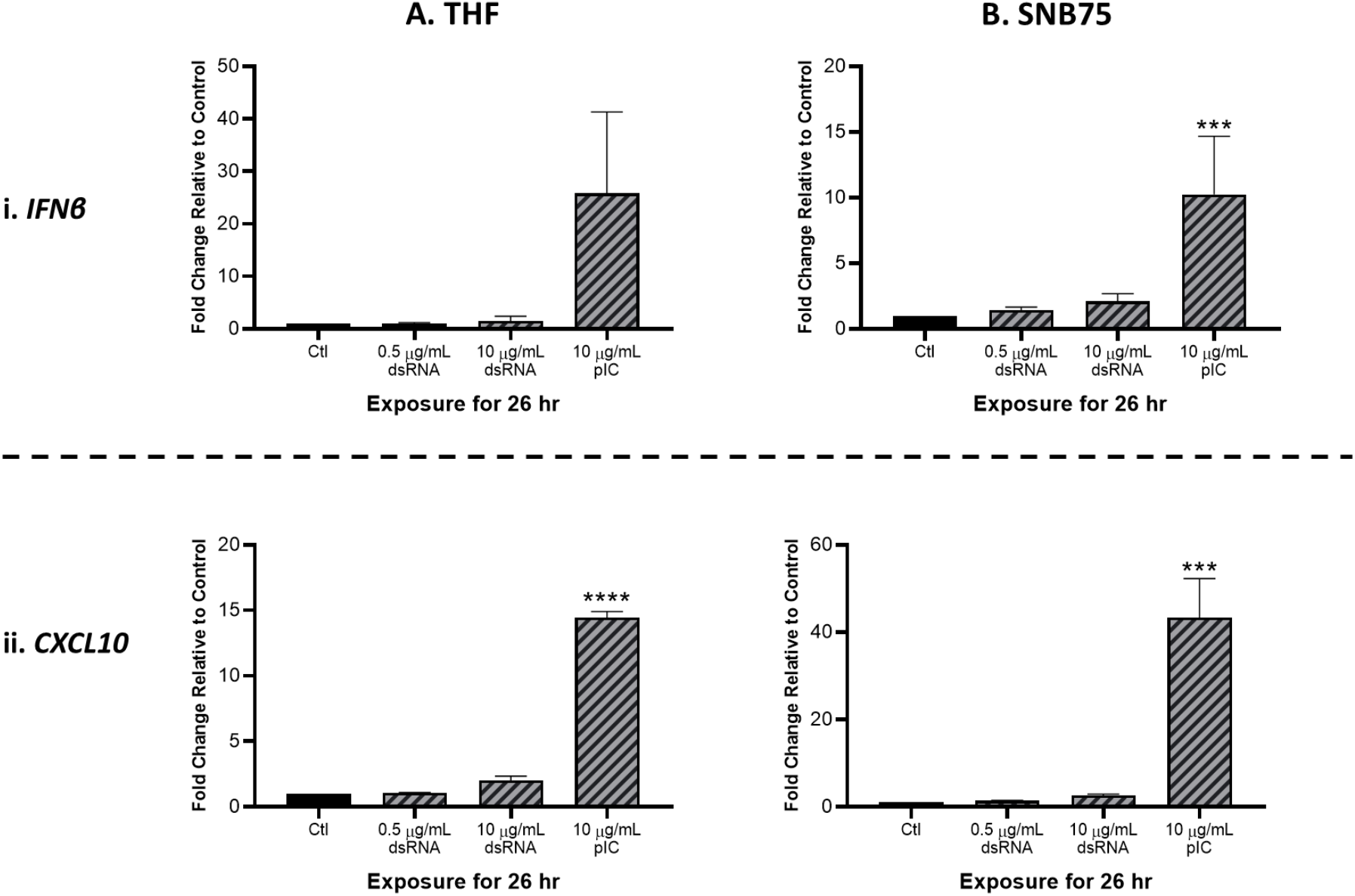
Type I IFN gene expression is not stimulated by soaking cells with low doses of dsRNA. Both THF (**A**) and SNB75 (**B**) cells were soaked with 700 bp dsRNA for 26h at concentrations of 0.5 μg/mL and 10 μg/mL as well as with HMW pIC at a concentration of 10 μg/mL. Following treatment, transcript expression of *IFNβ* (**i**) and *CXCL10* (**ii**) was assessed via qRT-PCR analysis. All data were normalized to the reference gene (*β-Actin*) and expressed as a fold change over the control group where control expression was set to 1. Error bars represent +SEM, and represents the average of 3 independent replicates. A p-value of less than 0.001 is represented by a *** symbol while a p-value of less than 0.0001 is represented by a **** symbol when compared only to the control (Ctl) treatment.

### 3.2 Soaking with long dsRNA does not negatively impact the viability of mammalian cells

To validate that soaking with long dsRNA does not negatively influence the health status of THF, SNB75 or MRC5, cell survival and metabolism were both measured. Following 24h exposure to a range of 700 bp GFP dsRNA concentrations, cellular metabolism was shown to significantly increase at only the highest dsRNA concentration assessed, 800 ng/mL (**Figure 2A**). The toxicity experiments did not use dsRNA concentrations greater than 800 ng/mL because higher concentrations were unnecessary to see RNAi effects. The significant increase in cellular metabolism at 800 ng/mL was observed in THF (**Figure 2Ai**), SNB75 (**Figure 2Aii**) and MRC5 (**Figure 2Aiii**) when compared to the 0 ng/mL control. Meanwhile, membrane integrity was shown to not be influenced at any of the dsRNA concentrations assessed in all three of the cell lines studied (**Figure 2B**). None of the dsRNA treated cells presented values significantly lower than the control cells, for both Alamar Blue and CFDA, indicating none of the dsRNAs treated were cytotoxic.

**Figure 2:**
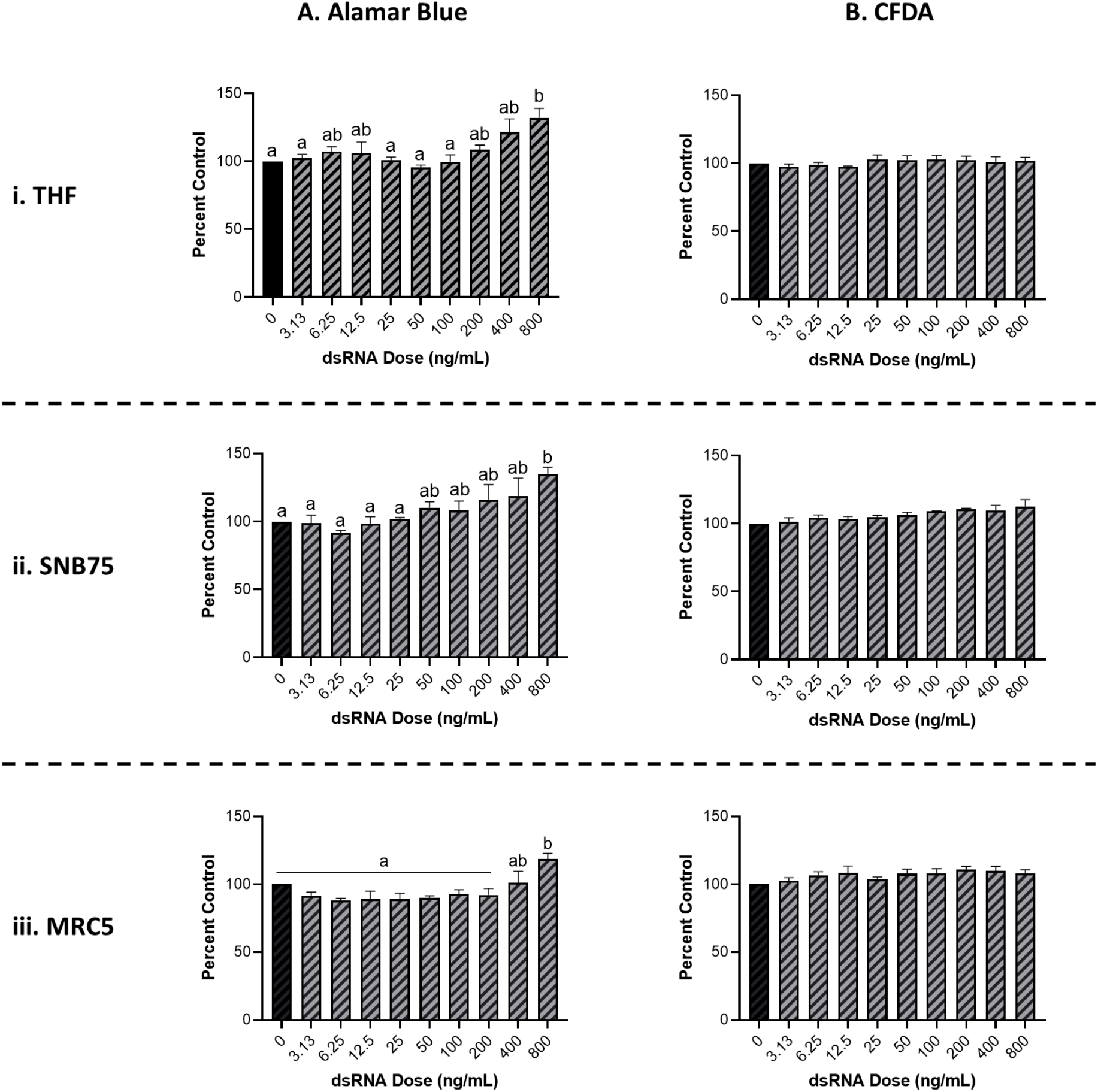
Soaking cells with long dsRNA does not negatively influence cell viability. THF (**i**), SNB75 (**ii**) and MRC5 (**iii**) were soaked with 700 bp dsRNA for 26h at concentrations that ranged from 0 ng/mL to 800 ng/mL. Cellular metabolism was measured using an Alamar Blue assay (**A**) and membrane integrity was measured using CFDA (**B**). Error bars represent +SEM, and each data point represents the average of 3 independent experiments. A p-value of less than 0.05 was considered to be statistically significant. Error bars with different letters represent significantly different data.

### 3.3 Long dsRNAi can only be stimulated by dsRNA lengths of 400 bp or greater

It was initially observed that pre-soaking cells with 500 ng/mL of 700 bp GFP dsRNA for 2h could stimulate protection towards VSV-GFP in both THF and SNB75 (**Figure 3**). Cell viability (**Figure 2**) and IFN induction by dsRNA (**Figure 1**) were both measured using 700bp long GFP dsRNA; however, the length of dsRNA capable of inducing dsRNAi required optimization. It was observed that dsRNA of 300 bp and shorter could not significantly induce knockdown of VSV-GFP in THF cells (**Figure 3A**). In the cancerous SNB75 cell line, the dsRNA length cut-off was less definitive as both 300 and 400 bp did not significantly differ from either the control condition or those inducing significant knockdown (**Figure 3B**). The appearance of the VSV-GFP infected THF following the dsRNA treatments revealed whether knockdown was occurring as the level of fluorescence is directly related to viral load (**Figure 3C**).

**Figure 3:**
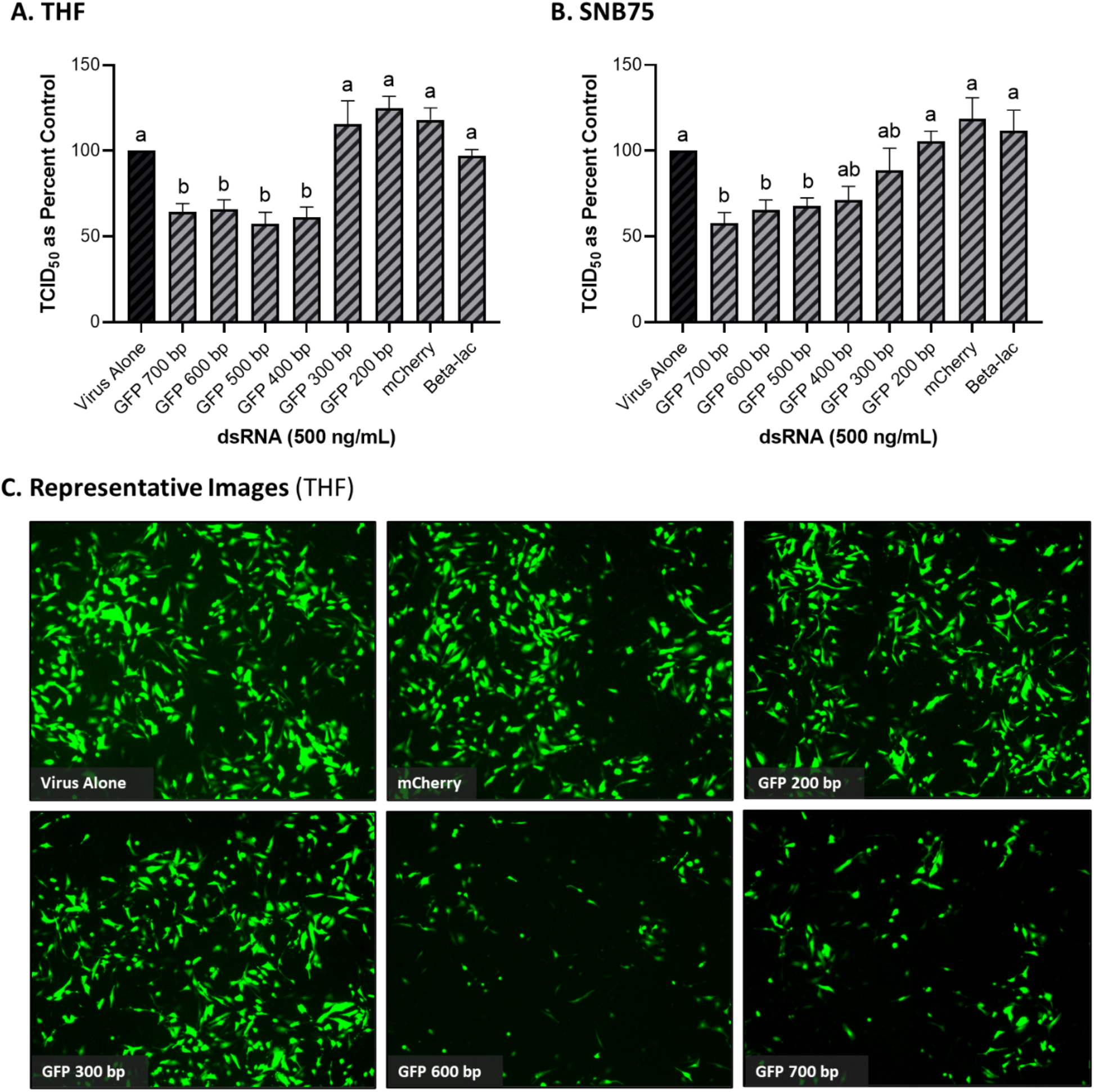
Viral knockdown is observed when pre-soaking cells with sequence-specific long dsRNA and this response is length dependent. THF (**A**) and SNB75 (**B**) cells were pre-soaked with either sequence specific (GFP) dsRNA ranging from 200 bp to 700 bp in length, non-sequence specific (mCherry or beta-lac) dsRNA of 700 bp or DPBS as a control for 2h prior to 24h infection with VSV-GFP (MOI = 0.1). Appearance of the THF cells after treatments with dsRNA and VSV-GFP infection as observed under the fluorescent microscope at 50X magnification (**C**). Error bars represent +SEM, and each data point represents the average of 6 independent replicates. A p-value of less than 0.05 was considered to be statistically significant. Error bars with different letters represent significantly different data.

### 3.4 Mammalian cells soaked with long dsRNA of viral genes can induce viral knockdown

Because GFP is not a naturally occurring gene found in viruses, the ability of dsRNA encoding viral gene sequences to stimulate dsRNAi was explored next. Soaking mammalian cells with viral gene specific dsRNA was shown to induce knockdown of corresponding viruses (**Figure 4**). When THF and SNB75 were pre-soaked with 500 ng/mL of 700 bp dsRNA synthesized to the N and M protein genes of VSV, significant knockdown was observed when compared to the non-sequence matched controls of mCherry and Beta-lac (**Figure 4A** and **4B**). Additionally, when a mixture of 250 ng/mL N protein dsRNA and 250 ng/mL M protein dsRNA was used to pre-soak THF cells, significant knockdown was still observed but was comparable to when only 500 ng/mL of either dsRNA was used (**Figure 4A**). For SNB75, this mixture pre-exposure was not observed to be significantly different to the control (**Figure 4B**). When MRC5 cells were pre-soaked with 500 ng/mL of 700 bp dsRNA synthesized to the RdRp, N protein, M protein and spike protein genes of HCoV-229E, significant reduction of viral particle production was observed for all exposures except for the RdRp dsRNA (**Figure 4C**). In Calu-3 cells, significant reduction in viral replication was observed after pre-treatment with 1000 ng/mL of SARS-CoV-2 N protein dsRNA when compared to both the virus alone control and the mis-matched mCherry dsRNA control (**Figure 4D**). However, no significant viral inhibition was observed when Calu-3 cells were pre-treated with 1000 ng/mL of SARS-CoV-2 M protein dsRNA (**Figure 4D**).

**Figure 4:**
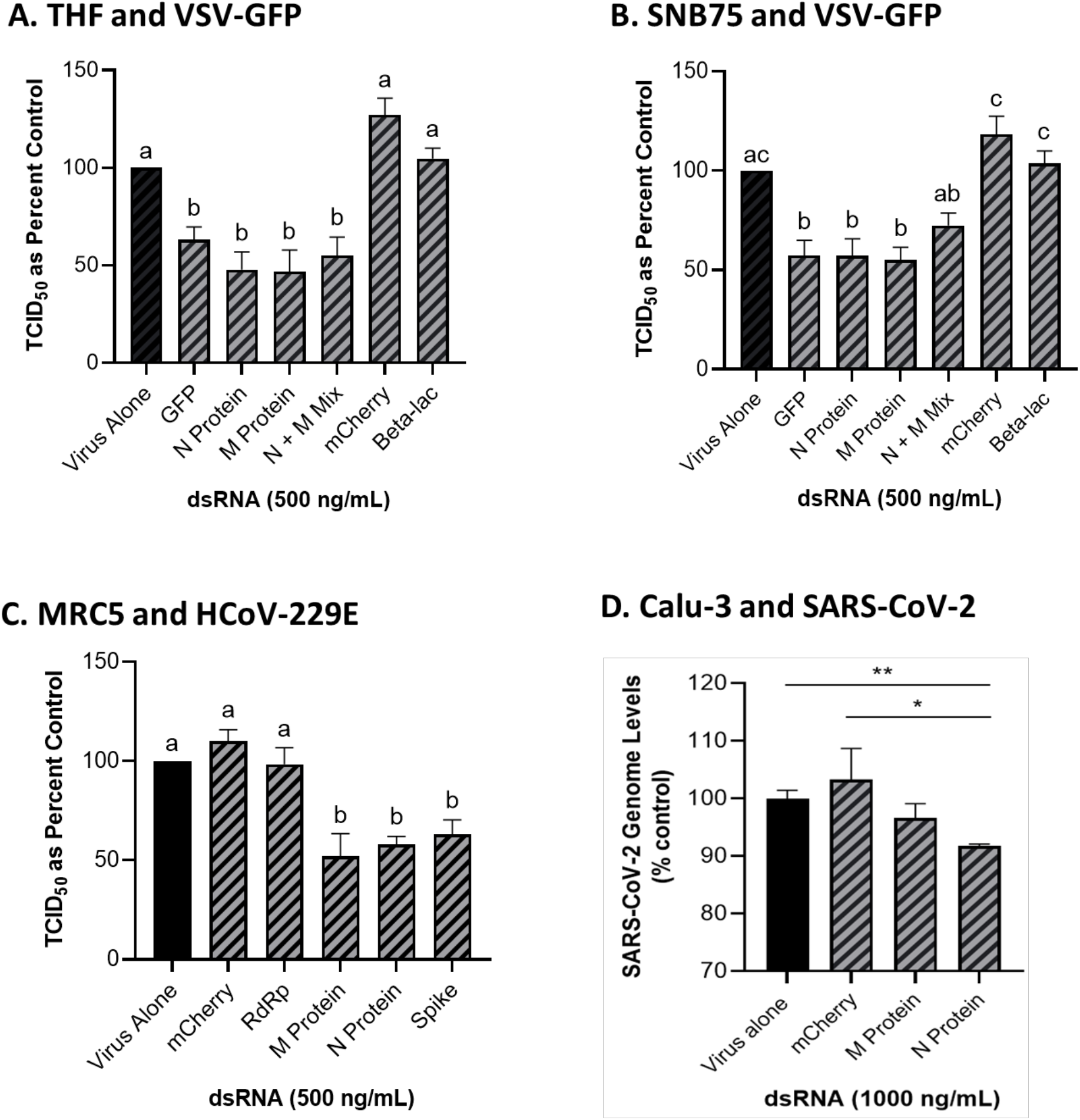
Soaking cells with long dsRNA of viral genes can induce knockdown of the complementary virus. Both THF (**A**) and SNB75 (**B**) cells were pre-soaked for 2h with either DPBS alone, 500 ng/mL of the mis-matched dsRNA controls (mCherry or Beta-lac), 500 ng/mL of VSV N protein dsRNA, 500 ng/mL of VSV M protein dsRNA or a mixture of 250 ng/mL N protein dsRNA with 250 ng/mL of M protein dsRNA before infection with VSV-GFP (MOI = 0.1) for 24h. MRC5 cells were pre-soaked for 2h with either DPBS alone, 500 ng/mL of the mCherry mis-matched dsRNA sequence control or 500 ng/mL of dsRNA matching HCoV-229E sequences for either RdRp, M protein, N protein and the spike protein before 24h infection with HCoV-229E (MOI = 0.02) (**C**). Calu-3 cells were pre-soaked for 2h with either DPBS alone, 1000 ng/mL of the mCherry mis-matched dsRNA sequence control or 1000 ng/mL of dsRNA matching SARS-CoV-2 sequences for either M protein and N protein prior to 24h infection with SARS-CoV-2 (MOI = 1.0) (**D**). Error bars represent +SEM, and each data point represents the average of 6 independent replicates. A p-value of less than 0.05 was considered to be statistically significant and different letters represent significant differences. For the SARS-CoV-2 data, a p-value of less than 0.01 is represented by a ** symbol and a p-value of less than 0.05 is represented by a * symbol when compared only to the control treatment

### 3.5 Knockdown via dsRNA soaking is also observed in human pBECs

In addition to the immortalized cell lines described above, the knockdown capability of pre-soaking cells with long dsRNA was also explored in primary pBEC cultures (**Figure 5**). An image of the pBECs after growth in culture for 28 days (**Figure 5A**). Significant knockdown of VSV-GFP was observed when pBECs were pre-treated with 500 ng/mL of dsRNA to the N protein of the virus when compared to the unmatched mCherry control (**Figure 5B**). Similar viral knockdown was also observed when the pBECs were pre-soaked with HCoV-229E M protein dsRNA which resulted in significant knockdown of HCoV-229E when compared to the mCherry control (**Figure 5C**). As a comparison it was also shown that soaking pBECs with 50 μg/mL of pIC also induced antiviral protection (**Figure 5C**). Indeed the level of protection provided by pIC was comparable to that provided by M protein encoding dsRNA (*p* = 0.1053442).

**Figure 5:**
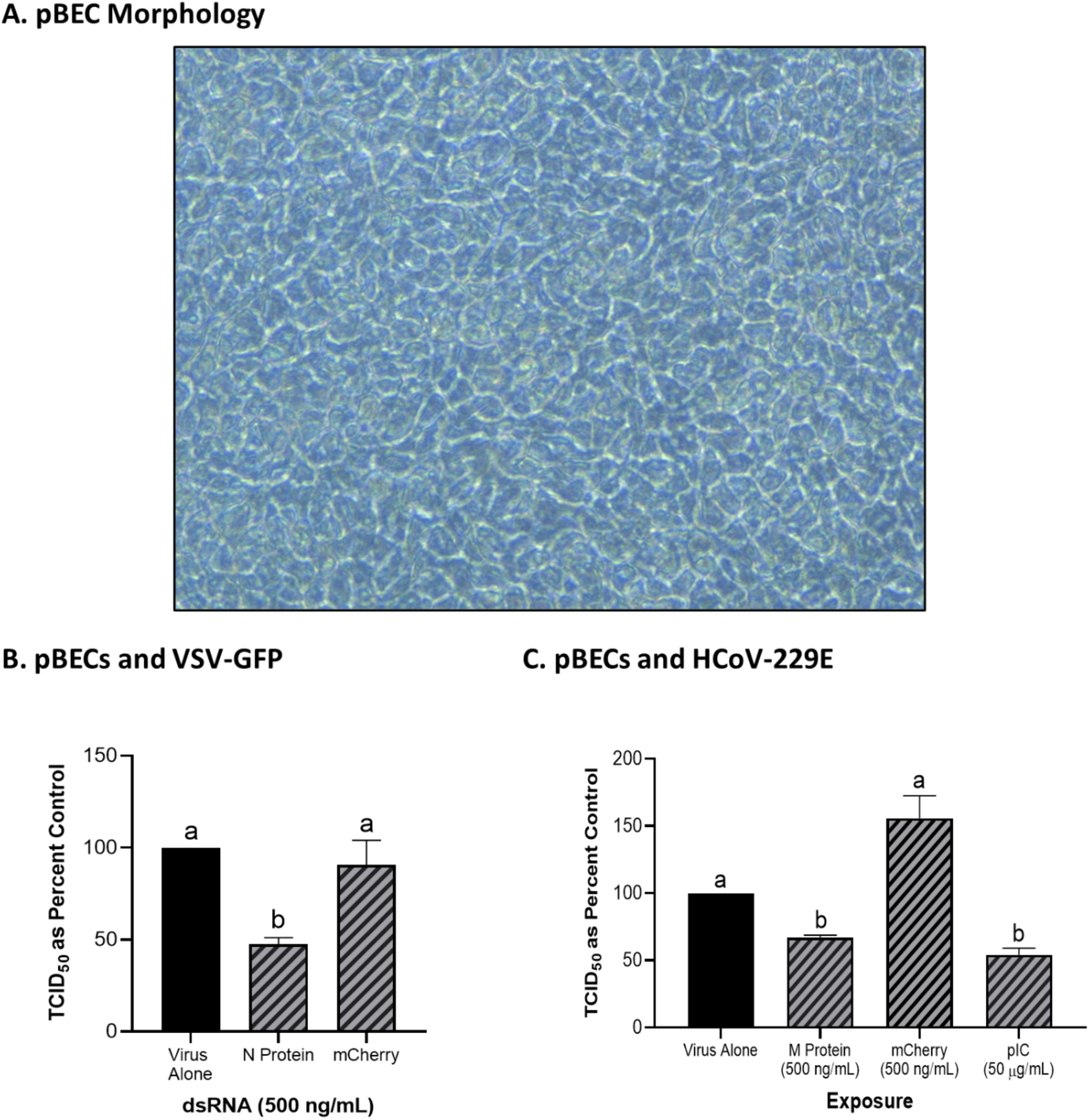
Primary Bronchial Epithelial/Tracheal Cells (pBECs) pre-soaked with long dsRNA of viral genes inhibits infection with corresponding viruses. The pBECs were grown to confluence and were shown to exhibit characteristics indicative of epithelial/tracheal cells, including mucous production and cilia function (**A**). The pBECs were pre-soaked with either DPBS, 500 ng/mL of the mis-matched mCherry dsRNA control or 500 ng/mL of VSV N protein dsRNA before infection with VSV-GFP (MOI = 0.1) for 24h (**B**). The pBECs were also pre-treated with either DPBS, 50 μg/mL of HMW pIC, 500 ng/mL of the mis-matched mCherry dsRNA control or 500 ng/mL of HCoV-229E M protein dsRNA before infection with HCoV-229E (MOI = 0.1) for 24h (**C**). Error bars represent +SEM, and each data point represents the average of 3 independent replicates. A p-value of less than 0.05 was considered to be statistically significant. Error bars with different letters represent significantly different data.

### 3.6 Soaking cells with siRNA did not induce viral knockdown

When both THF and SNB75 cells were pre-soaked with GFP siRNA, TCID_50_ levels of VSV-GFP were comparable to the unstimulated control and to the mis-matched long dsRNA mCherry control (**Figure 6Ai** and **6Bi**). Meanwhile, soaking these cells with 700 bp GFP dsRNA was again shown to induce significant knockdown of the VSV-GFP virus (**Figure 6Ai** and **6Bi**). This result was not due to inefficacy of the siRNA molecules as transfecting THF and SNB75 with the GFP siRNA induced significant knockdown when compared to transfection with the negative control siRNA (**Figure 6Aii** and **6Bii**). At the timepoint tested the knockdown of VSV-GFP by GFP siRNA is similar to that by 700 bp GFP dsRNA.

**Figure 6:**
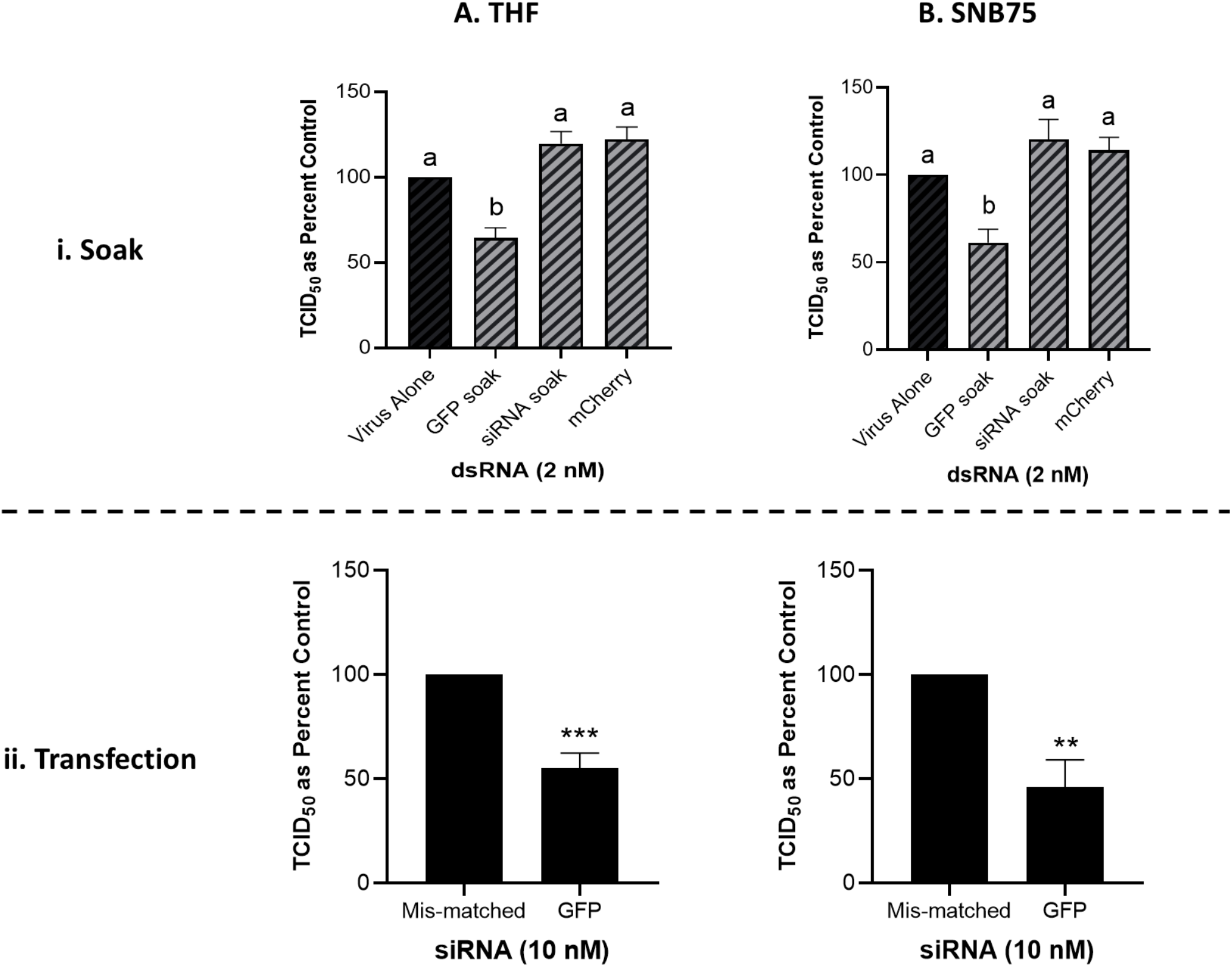
Soaking is sufficient for long dsRNA-induced antviral effects but not siRNA. THF (**Ai**) and SNB75 (**Bi**) were pre-soaked for 2h with either DPBS, 2 nM of long mCherry dsRNA, 2 nM of long GFP dsRNA or 2 nM of GFP siRNA prior to infection with VSV-GFP (MOI = 0.1) for 24h. To ensure that the siRNA was functional, THF (**Aii**) and SNB75 (**Bii**) cells were transfected with either 10 nM of GFP siRNA or 10 nM of the negative control siRNA for 24h prior to infection with VSV-GFP (MOI = 0.1) for 24h. Error bars represent +SEM, and each data point represents the average of 5 independent replicates. A p-value of less than 0.05 was considered to be statistically significant and different letters represent significant differences. For the transfection data, a p-value of less than 0.01 is represented by a ** symbol while less than 0.001 is represented by a *** symbol.

### 3.7 Long, synthetic combination dsRNA molecules can inhibit VSV-GFP via multiple gene knockdown

Combination dsRNA molecules were synthesized to test whether multiple VSV genes could be knocked down when 700 bp of dsRNA contained sequences for two different viral genes. **Figure 7A** is a schematic of the three different combination dsRNA molecules that were synthesized for this study. When THF cells were pre-treated with 500 ng/mL of each combination dsRNA molecule and the mCherry unmatched sequence control, only the three combination molecules were able to induce significant knockdown of VSV-GFP (**Figure 7Bi**). When measuring gene expression, only the 5’N-3’M molecule was able to induce significant knockdown of both the VSV N protein and M protein genes (**Figure 7Bii** and **7Biii**). When THF cells were pre-exposed to 1000 ng/mL of the combination dsRNA molecules and the mCherry control, it was observed again that only the three combination molecules induced significant knockdown of VSV-GFP (**Figure 7Ci**). Through pre-soaking with 1000 ng/mL, only the 5’N-3’M molecule induced significant knockdown of the VSV N protein gene (**Figure 7Cii**), but both 5’N-3’M and the N-M Alt molecules were able to induce significant knockdown of the VSV M protein gene (**Figure Ciii**).

**Figure 7:**
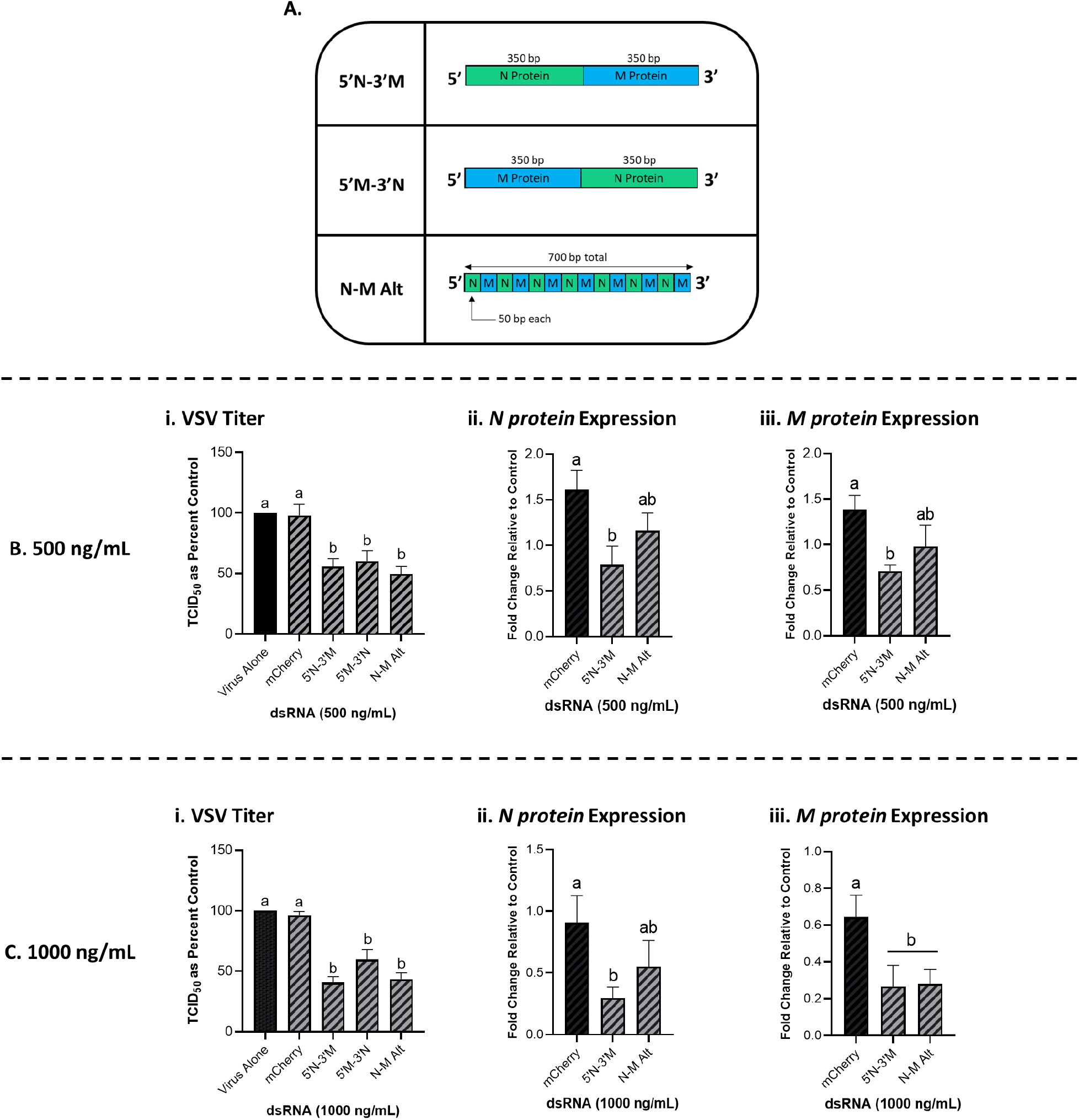
Combination dsRNA molecules can inhibit viruses through the knockdown of multiple viral genes. Three different 700 bp combination genes were synthesized using gBlocks referred to as 5’N-3’M, 5’M-3’N and N-M Alt (**A**). THF cells were pre-soaked for 2h with DPBS or 500 ng/mL of either mCherry, 5’N-3’M, 5’M-3’N or N-M Alt before being exposed to VSV-GFP (MOI = 0.1) for 24h (**Bi**). Following this treatment, cells were collected and RNA extracted so that gene expression of the VSV N protein gene (**Bii**) and M protein gene (**Biii**) could be measured by qRT-PCR. THF cells were also pre-soaked for 2h with DPBS or 1000 ng/mL of either mCherry, 5’N-3’M, 5’M-3’N or N-M Alt before being exposed to VSV-GFP (MOI = 0.1) for 24h (**Ci**). Following this treatment, cells were collected and RNA extracted so that gene expression of the VSV N protein gene (**Cii**) and M protein gene (**Ciii**) could be measured by qRT-PCR. Error bars represent +SEM. Each data point for the titer data represents the average of 6 independent replicates while the qRT-PCR data represents the average of 5 independent replicates. A p-value of less than 0.05 was considered to be statistically significant. Error bars with different letters represent significantly different data.

### 3.8 Dicer1 is a required component for the viral knockdown stimulated via cell soaking with dsRNA

Knockdown of VSV-GFP was also obtained in the mouse MSC cell line that contains functional Dicer1 when pre-soaked with long dsRNA containing N protein sequence for 2h prior to infection (**Figure 8A**). In comparison, when using the matching cell line that was Dicer1-defective, the significant decrease in viral knockdown was abolished (**Figure 8B**).

**Figure 8:**
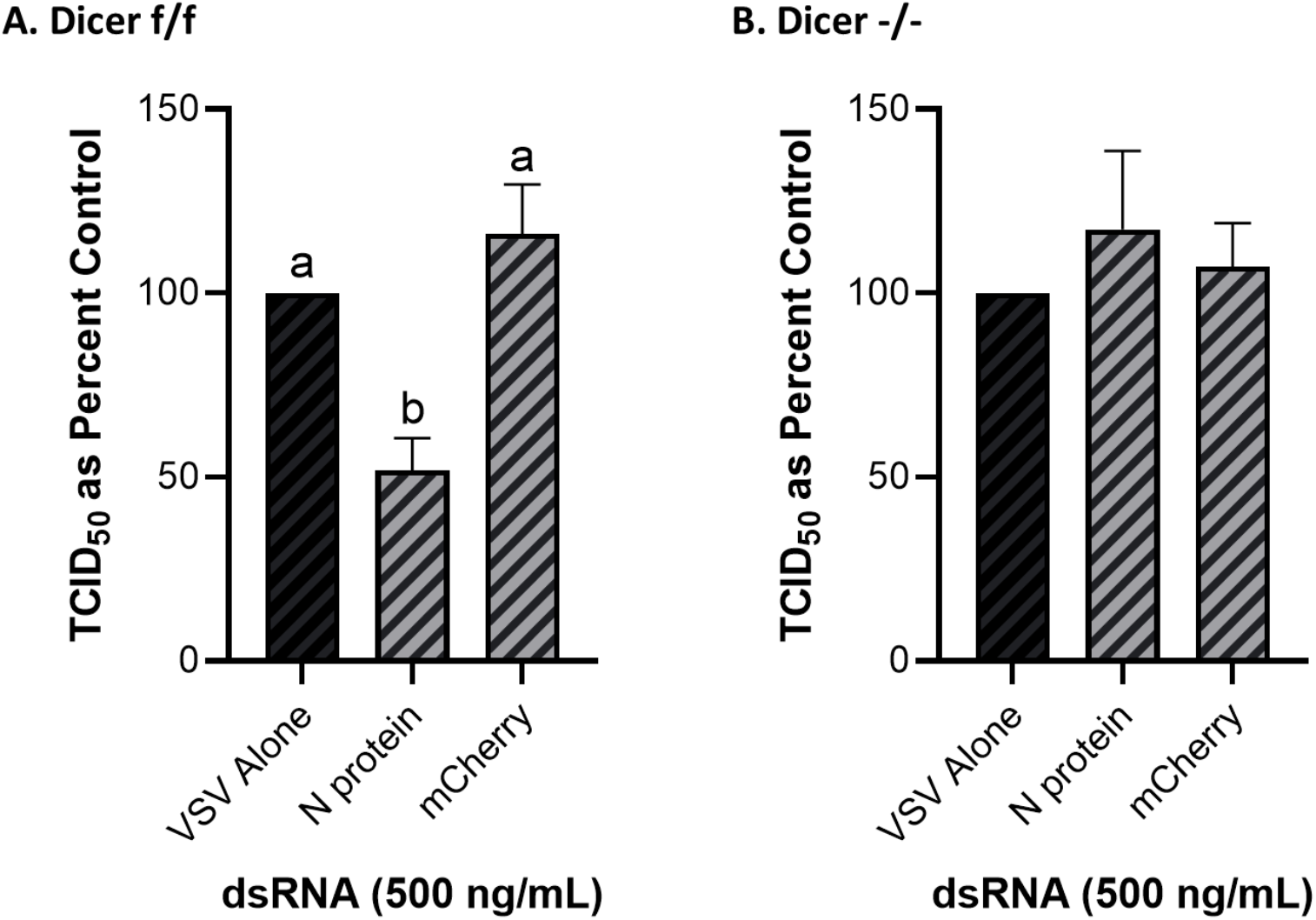
The observed viral inhibition by long dsRNA soaking is dependent on the presence of functional Dicer proteins. Mouse MSCs that had functional Dicer1 were pre-soaked for 2h with DPBS or 500 ng/mL of either mis-matched mCherry dsRNA or matched GFP dsRNA before infection with VSV-GFP (MOI = 0.1) for 24h (**A**). The matching mouse MSC cell line that was a KO for Dicer1 were also pre-soaked for 2h with DPBS or 500 ng/mL of either mis-matched mCherry dsRNA or matched GFP dsRNA before infection with VSV-GFP (MOI = 0.1) for 24h (**B**). Error bars represent +SEM, and each data point represents the average of 6 independent replicates. A p-value of less than 0.05 was considered to be statistically significant. Error bars with different letters represent significantly different data.

## 3. Discussion

It is well established that transfecting and/or soaking vertebrate cells with long dsRNA or pIC, the IFN pathway will be stimulated (Alexopoulu *et al*., 2001; Hemmi *et al*., 2004; DeWitte-Orr *et al*., 2009; De Waele *et al*., 2017). However, little to no research has explored what the lower limit is for stimulating this IFN response through cell soaking. One study by Hägele and colleagues (2009) revealed that soaking murine cells with low concentrations of pIC (0.1-3 μg/mL) did not stimulate protein production of IFNβ and CXCL10. However, when the cells were transfected with these low doses of pIC, a significant increase was observed for both IFNβ and CXCL10 protein production (Hägele *et al*., 2009). In the present study, we soaked human cells with 10 μg/mL of pIC and were successfully able to stimulate the gene expression of IFNβ and the ISG, CXCL10. Furthermore, we observed that the long dsRNA concentrations used in the present study did not induce the expression of these genes in both THF and SNB75. For the fibroblast cells used, IFNβ would be anticipated to be the primary IFN produced in response to dsRNA (Li *et al*., 2018). However, in glioblastoma cells, variations in IFN competence have been reported (Dick and Hubbell, 1987; Imaizumi *et al*., 2014; De Waele *et al.*, 2021). When previously explored in multiple glioblastoma cell lines, pIC was shown to modestly induce IFNβ expression but significantly induced ISG (ISG15 and CXCL10) expression in some of these cultures (Wollmann *et al*, 2007). When taken together, the results presented here for IFN stimulation via cell soaking are comparable to what has been reported previously in the literature.

Numerous studies have shown that exposing mammalian cells to dsRNA, pIC and/or viral infection hinders cellular metabolism (reviewed by Nellimarla and Mossman, 2014). However, most of these studies explore the impact of IFN-inducing concentrations, which would be expected to reduce metabolism through induction of the antiviral state (reviewed by Fritsch and Weichart, 2016). The results presented here provide evidence that soaking cells with concentrations of dsRNA that are too low to induce IFN does influence cellular metabolism. Surprisingly, as this concentration increases (while still being too low to induce IFN), we observed a significant, *increase* in cellular metabolism. This could be due to low level stimulation of PAMPs by dsRNA, which has been previously observed to enhance the metabolism of immune cells (human dendritic cells)(Everts *et al*., 2014). The non-immune cells used in the present study can be stimulated by PAMPs and act as important sentinels for microbial infections, including those of viral origin (reviewed by Bautista-Hernández *et al*., 2017). Thus, soaking cells with concentrations of dsRNA that do not induce IFN can stimulate metabolic rate through activation of IFN-independent innate antiviral processes within the cell.

Unless transfection is used, the literature supports that the RNAi pathway can only be induced if the original dsRNA molecule meets a certain length requirement. This length dependence was observed in the current study but has also been shown previously in various invertebrate models. The impact of dsRNA length on RNAi efficiency was recently explored in the Colorado potato beetle (He *et al*., 2020). Though the beetles were exposed through ingestion of dsRNA expressing potato plants rather than soaking, it was shown that 200 bp or greater was required to induce a robust RNAi response (He *et al*., 2020). A similar observation was observed when *Caenorhabditis elegans* was injected with long dsRNA. The length requirement for efficient knockdown was smaller in this example, at 50 to 100 bp (Parrish *et al*., 2000). When explored further, it was revealed that the minimal length of dsRNA required for efficient RNA uptake by *C. elegans* SID-1 is 50 bp (Feinberg and Hunter, 2003; Li *et al.*, 2015). Importantly, increasing the length of the dsRNA molecules has been shown to enhance the observed knockdown through RNAi. When soaking *Drosophila* S2 cells with 700 bp dsRNA, 95-99% knockdown of the target protein was observed (Clemens *et al*., 2000). Further study with S2 cells revealed that there was a clear length-dependence when soaking the cells with luciferase dsRNA that was not observed when transfecting them (Saleh *et al*., 2006). Though significant knockdown was still observed when soaking with shorter lengths, 200 bp and greater were found to be much more effective at inducing luciferase knockdown (Saleh *et al.*, 2006). Because there is no concern of stimulating the IFN response in invertebrate cells, concentration may also play a role that cannot be explored in IFN-competent mammalian cells. This may provide an explanation as to why the length requirement (~300-400 bp and greater) observed in the present study was greater than those described using invertebrate cells. It is also possible that the size specificity of SID-1 includes smaller dsRNA molecules when compared to SIDT2. Additionally, because the SIDT2 channel has a higher binding affinity for dsRNA lengths ranging from 300-700 bp (Li *et al*., 2015), this also supports its involvement here wherein knockdown was only achievable in THF and SNB75 using dsRNA lengths of 300-400 bp and greater.

Inhibition of viral infection through pre-stimulation of the RNAi pathway is not a novel concept and has been deemed successful against multiple mammalian viruses (Gitlin *et al*., 2002; Wheeler *et al*., 2013). In fact, higher efficiency has been reported when using siRNAs that are specific for certain viral genes over others, similarly to what was observed in the present study when using long dsRNA. When mammalian MDCK cells were pre-transfected for 8h with siRNA matching influenza viral genes of NP (nucleocapsid) and PA (component of RNA transcriptase), greater viral inhibition was observed when compared to siRNA developed for the genes of the M (matrix) and certain PB1complexes (component of RNA transcriptase, Ge *et al*., 2003). Moreover, when the same siRNAs were used in chicken embryos, only those that were very effective in the MDCK cells had protective effects *in vivo* (Ge *et al*., 2003). When exploring the use of siRNA for combatting COVID-19, Wu and Luo (2021) reported 50% inhibition rates in 24h when targeting the structural Spike, N and M protein genes of SARS-CoV-2 that were overexpressed in human epithelial cells. These results have also been replicated in live animal trials. In an *in vivo* trial, mice were injected with lentiviruses containing siRNA that targeted either the L (polymerase) or N (nucleocapsid) protein of the rabies virus (RV). It was found that targeting the structural N protein provided 62% protection to RV infected mice while no protection was observed when the L protein was the target (Singh *et al*., 2014). Based on these previous results, it appears that the type of virus, and likely, variances in replication processes, play a role in which target genes have higher efficiency for RNAi knockdown. When exploring a rhabdovirus and two coronaviruses in the current study, pre-soaking with long dsRNA matching structural genes (N, M and spike proteins) was observed to be more successful than those associated with the viral transcriptional machinery (RdRp). A systematic study of each gene, including sequences within each gene, is needed in future studies to better understand what sequences are optimal targets for suppressing virus replication via dsRNAi.

As treatments that stimulate dsRNAi towards a single viral gene were successful, so simultaneous inhibition of multiple viral genes would be anticipated to enhance this effect. Combination treatments with siRNAs have shown promise in the suppression of various viral pathogens. When siRNAs that targeted both the G (glycoprotein) and the N protein genes of rabies virus was expressed in mammalian cells using a single cassette, an 87% reduction of the target virus was observed (Meshram *et al*., 2013). It should be noted that individual sequences offered an 85% reduction in virus titres. Similarly, when rat fibroblast cells were exposed to combination siRNAs targeting both the Immediate-early-2 and DNA polymerase genes, a significant reduction in associated mRNAs and cytopathic effects was observed following infection with a novel rat Cytomegalovirus (Balakrishnan *et al*., 2020). This siRNA combination inhibition has also been explored *in vivo* using both rhesus monkeys and macaques. SiRNA combinations targeting multiple genes of the Zaire Ebola virus (ZEBOV) provided 66% protection in the rhesus monkeys and 100% protection in macaques to lethal doses of ZEBOV when this treatment was administered in stable nucleic acid lipid particles (Geisbert *et al*., 2010). Due to the greater length of long dsRNA when compared to their siRNA counterparts, it is possible to have multiple viral genes sequences present in a single molecule. In theory, this could induce knockdown of multiple viruses or multiple viral genes to inhibit infection, all without the requirement of transfection or creation of multiple dsRNA fragments. When this was explored for the first time in the present study, three combination molecules for the VSV N and M protein were shown to significantly knockdown viral titers when cells were soaked with the long dsRNA. However, qRT-PCR analysis revealed that only one of these molecules (5 ‘N-3’M) was able to significantly reduce mRNA levels of both viral genes. Though mRNA degradation is often associated with the knockdown observed during RNAi, it is important to recognize that the RISC complex can bind to complimentary mRNAs and in doing so, repress translation (reviewed by van den Berg *et al*., 2008). As a result, mRNA expression may not decrease but the associated protein levels would be reduced (Alemán *et al*., 2007; Ma *et al*., 2013). This provides an explanation as to why viral titers decreased, but mRNA levels were not always significantly reduced.

One of the defining mechanisms within the RNAi pathway is the cleavage of long dsRNAs into siRNAs by Dicer proteins. In both vertebrates and invertebrates, functional Dicer has been shown to be a necessity for the sequence-specific knockdown associated with RNAi (Bernstein *et al*., 2001; Ketting *et al*., 2001; Zhang *et al.*, 2002; Sakurai *et al.*, 2011). We confirmed this in mammalian cells as only mouse MSCs with functional Dicer were able to induce significant knockdown of viral titers when pre-soaked with long, sequence-matched dsRNA. The role of Dicer in inhibiting viral infection has been explored in mammalian cells, but viral replication has only been modestly affected in Dicer knockouts (Matskevich and Moelling, 2007; Bogerd *et al*., 2014). Notably, these cells were not pre-treated with sequence-specific dsRNA prior to these infections. Based on the results of the present study, it appears that pre-soaking cells with low doses of dsRNA can provide sequence-matched protection against complimentary viral pathogens. Perhaps cells will default to RNAi when viral levels are not high enough to stimulate the IFN pathway. Aside from the data presented, this is also supported by evidence from numerous viral pathogens that specifically inhibit various components of the RNAi pathway, including Dicer. (Wang *et al*., 2006; Qui *et al*., 2020). To successfully establish infection when low levels of virus particles are present within the cell, perhaps it is critical for viruses to overcome the initial antiviral response via dsRNAi. This initial RNAi based disruption of viral mRNA could represent a constant, sentinel-like antiviral mechanism in mammalian cells. In this context mammalian cells would mount an RNAi based inhibition of viral mRNA at lower levels of circulating viral dsRNA, without initiating the energy consuming IFN response.

The results of the present study indicate several unique findings. Firstly, this is the first time in mammalian cells that RNAi has been observed through the natural uptake of sequence-specific dsRNA. This indicates that it may be possible to develop antiviral therapies involving long dsRNAs that do not involve costly transfection agents or stimulation of the damaging IFN response. Second, this viral inhibition was observed to be length-dependent, as only dsRNA that was 300-400 bp in length or greater would induce knockdown. This strongly implies a molecular channel such as SIDT2, although this was not explicitly confirmed in our study. Finally, the success of combination dsRNA constructs suggest that it may be possible to target either multiple genes within a single virus, genes originating from more than one virus or possibly those from one virus along with associated host proteins. Moreover, we were able to provide evidence that the observed viral inhibition was due to RNAi as Dicer1 knockouts could not induce this response. Confirming that pre-stimulation of RNAi in mammalian cells will induce protection against a variety of viral pathogens could have important implications for the development of novel antiviral therapies.

## 4. Materials and Methods

### 4.1 Immortal Cell Maintenance

The telomerase-immortalized human fibroblast cell line, THF, was received as a generous gift from Dr. Victor DeFilippis of Oregon Health and Science University. The THF cells were maintained in Dulbecco’s modified eagle medium (DMEM, Corning). The human glioblastoma cell line, SNB75, was obtained as part of the NCI-60 panel. The SNB75 cells were cultured in Roswell Park Memorial Institute (RPMI)-1640 medium (Corning). Two human embryonic lung cell lines, MRC-5 (CCL-171) and HEL-299 (CCL-137) were obtained from the American Type Culture Collection (ATCC). The MRC-5 cells were propagated in Eagle’s Minimum Essential Medium (EMEM, Corning) while the HEL-299 cells were cultured in DMEM (Corning). The Dicer1 positive murine mesenchymal stem cell (MSC) line (Dicer1 f/f, CRL-3220) and the Dicer1 KO MSC line (Dicer1 -/-, CRL-3221) were obtained from the ATCC. Both mouse cell lines were cultured in Minimum Essential Medium (MEM) Alpha (Gibco). The media used for the above cell lines were supplemented with 10% fetal bovine serum (FBS, Seradigm) and 1% penicillin-streptomycin (P/S, Sigma). The lung adenocarcinoma cell line, Calu-3 (HTB-55), and the green monkey kidney epithelial cell line, Vero E6 (CRL-1586), were also obtained from ATCC. Vero E6 cells were cultured in DMEM supplemented with 10% FBS, 1x L-glutamine and 1% p/s. The Calu-3 cells were regularly cultured in MEM Alpha medium (Corning) supplemented with 10% FBS, 1% P/S and 1% HEPES. All cell lines were grown in vented T75 flasks (Falcon) at 37°C with 5% CO_2_.

### 4.2 Viruses

#### 4.2.1 Viral Propagation

The Indiana serotype of Vesicular Stomatitis Virus (VSV-GFP) contains a GFP gene incorporated between the viral G and L genes (Dalton and Rose, 2001). As a result, cells infected with VSV-GFP fluoresce green. In the present study, VSV-GFP was propagated on monolayers of HEL-299. Virus infections were performed in DMEM containing 10% FBS and 1% P/S at 37°C with 5% CO2. Virus-containing media was cleared of cellular debris by centrifugation at 4,000 g for 4 min followed by filtration through a 0.45 μm filter. The filtered supernatant was aliquoted and stored at −80°C until ready to be quantified.

HCoV-229E was purchased from ATCC (VR-740) and subsequently propagated on monolayers of MRC5 cells. Viral infections were performed in EMEM medium containing 2% FBS and 1% P/S at 37°C with 5% CO_2_. After two days, virus-containing media was cleared at 4,000 g and filtered through a 0.45 μM filter. The filtered supernatant was aliquoted and stored at −80°C until ready to be quantified. The second HCoV used in this study, SARS-CoV-2 isolate SB3, was propagated on Vero E6 cells as previously described by Banerjee *et al*. (2020). All SARS-CoV-2 infections were performed at a designated BSL-3 lab in accordance with guidelines from McMaster University.

#### 4.2.2 Tissue Culture Infectious Dose (TCID50)

The cell lines described above for viral propagation were seeded in 96-well plates (1.5 x 10^4^ cells/well) and were used to titer their respective viruses by TCID_50_. Following overnight adherence, all wells received 100 μL of fresh 10% FBS media. For VSV-GFP, the supernatants of interest were serially diluted 1:5 in basal DMEM media. With HCoV-229E, the supernatants of interest were serially diluted 1:10 in basal EMEM media. For each sample dilution, 10 μL was added to eight wells of a 96-well plate for VSV-GFP and six wells for HCoV-229E. Plates were then either incubated at 37°C (VSV-GFP) or 33°C (HCoV-229E) at 5% CO_2_. At three (VSV-GFP) or seven (HCoV-229E) days post-infection, wells were scored by the presence of cytopathic effects (CPE) and viral titers were calculated using the Reed and Meunsch method to obtain the TCID50/mL (Reed & Muensch, 1938).

### 4.3 Synthesis of dsRNA molecules

Genes of interest were amplified using forward and reverse primers that contained T7 promoters. The primer sets and their associated templates are outlined in **Table 1**. The DNA products were amplified by PCR using 10 ng of appropriate template (**Table 1**), 2X GOTaq colorless master mix (Promega), 0.5 μM of both forward and reverse primers (**Table 1**, Sigma Aldrich), and nuclease free water to a final volume of 50 μL. The following protocol was carried out in a Bio-Rad T100 thermocycler: 98°C - 5 min, 34 cycles of 98°C - 10s, 50°C - 10s, 72°C - 50s, followed by 72°C - 5min. The resulting DNA amplicons with T7 promoters on both DNA strands were purified using a QIAquick PCR purification kit (Qiagen). The purified product was then used in the MEGAScript RNAi Kit (Invitrogen) as per the manufacturer’s instructions to produce dsRNA. To confirm primer specificity, 100 ng of all PCR amplicons and the final dsRNA product were separated on 1% agarose gels containing 1% GelGreen (Biotium Inc.).

**Table 1:**
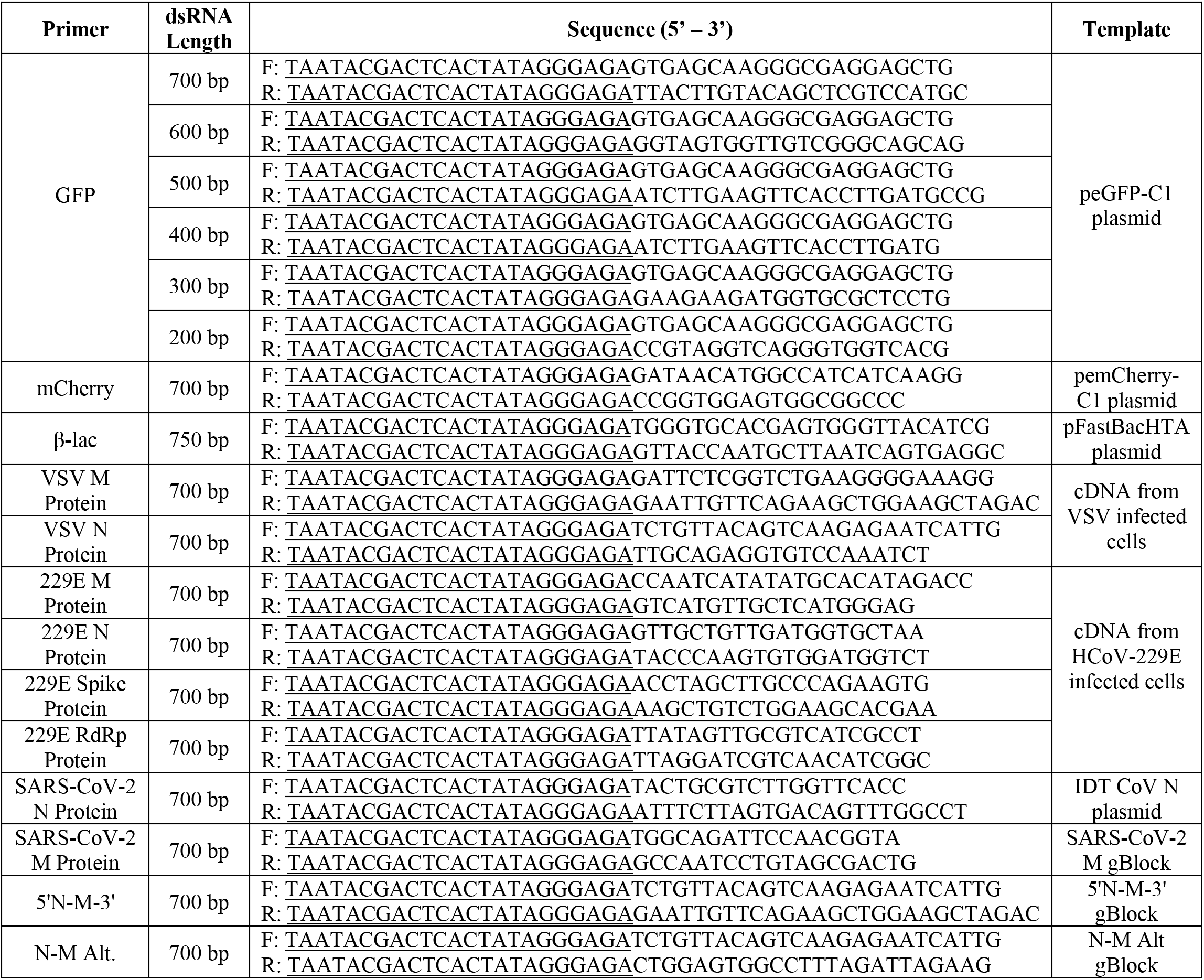
Primers with underlined T7 promoter sequences that were used for amplification of genes of interest for dsRNAi. The resulting DNA amplicons were then used for dsRNA synthesis. The resultant dsRNA length and the original template DNA used for each primer set is also outlined.

### 4.4 Testing for induction of the antiviral interferon response

#### 4.4.1 RNA Extraction

In a 24-well plate, either THF or SNB75 were seeded at a density of 5.0 x 10^4^ cells/well. Following overnight adherence, the media was replaced before exposure to either a DPBS control, 0.5 μg/mL of long dsRNA, 10 μg/mL of long dsRNA, or 10 μg/mL of high molecular weight (HMW) polyinosinic:polycytidylic acid (pIC) all diluted in full growth media. Cells were exposed to these treatments for 26h before the media was removed and the test wells were washed once with DPBS. Cells were then collected in TRIzol (Invitrogen) and total RNA was extracted according to the manufacturer’s instructions. RNA was then treated with Turbo DNA-free™ Kit (Invitrogen) to remove any contaminating genomic DNA. Complementary DNA (cDNA) was synthesized from 500 ng of purified RNA using the iScript™ cDNA Synthesis Kit (Bio-Rad) following protocols provided by the manufacturer.

#### 4.4.2 qRT-PCR

The expression of IFN related genes (IFNβ and CXCL10) was measured by quantitative real-time polymerase chain reaction (qRT-PCR). IFNβ was chosen because it is frequently the first type I IFN induced following dsRNA treatment, particularly in fibroblasts (Bolivar *et al*., 2018; Li *et al*., 2018). CXCL10 was chosen as the representative ISG because its expression levels are very high in the presence of IFNs (Buttmann *et al*., 2007; Antonelli *et al*., 2010). All PCR reactions contained: 2 μL of diluted cDNA, 2x SsoFast EvaGreen Supermix (Bio-Rad), 0.2 mM of forward primer (Sigma Aldrich), 0.2 mM of reverse primer (Sigma Aldrich) and nuclease-free water to a total volume of 10 μL (Fisher Scientific). The sequences and accession number for each primer set are outlined in **Table 2**. The qRT-PCR reactions were performed using the CFX Connect Real-Time PCR Detection System (Bio-Rad). The program used for all reactions was: 98°C denaturation for 3 min, followed by 40 cycles of 98°C for 5 sec, 55°C for 10 sec, and 95°C for 10 sec. A melting curve was completed from 65°C to 95°C with a read every 5 sec. Product specificity was determined through single PCR melting peaks. All qRT-PCR data was analyzed using the ΔΔCt method and is presented as the average of four experimental replicates with the standard error of the mean (SEM). Specifically, gene expression was normalized to the housekeeping gene (β-actin) and presented as fold changes over the control group.

**Table 2:**
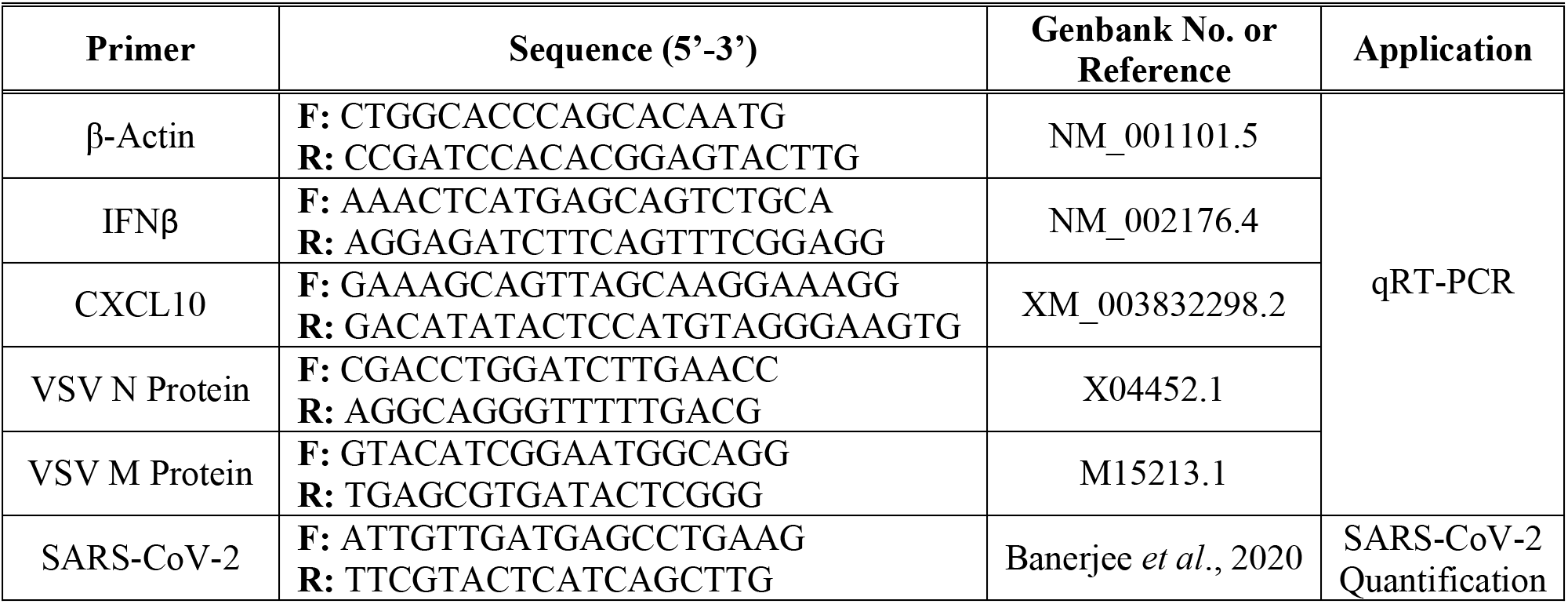
Primers used for qRT-PCR analyses and for SARS-CoV-2 qPCR quantification.

### 4.5 Cell viability to dsRNA

To determine whether different dsRNA concentrations could influence the survival of THF, MRC5 and SNB75, two fluorescent indicator dyes, Alamar Blue (AB, Invitrogen) and 5-carboxyfluorescein diacetate acetoxymethyl ester (CFDA-AM, Invitrogen), were used. Together these dyes provide an excellent indication of cell viability as both cellular metabolism (AB) and membrane integrity (CFDA-AM) are measured (Dayeh *et al*., 2003). THF and SNB75 cells were seeded at a density of 1 x 10^4^ cells/well in a 96-well tissue culture plate and allowed to adhere overnight at 37°C with 5% CO_2_. All cell monolayers were washed once with DPBS and then treated in eight-fold with a doubling dilution of dsRNA ranging from 800 ng/mL to 3.13 ng/mL for 24h at 37°C with 5% CO_2_ in normal growth media. Following incubation, each well was washed twice with DPBS before exposure to AB and CFDA-AM as described previously by Dayeh *et al.* (2003). Because two fluorescent dyes were used to test cell viability, the 96-well plate was read at an excitation of 530 nm and an emission of 590 nm for AB as well as an excitation of 485 nm and an emission of 528 nm for CFDA-AM. The reads were completed using a Synergy HT plate reader (BioTek Instruments). For each cell line analyzed, three independent experiments were performed.

### 4.6 Stimulating viral inhibition via soaking with low doses of dsRNA

THF and SNB75 cells were seeded at a density of 5.0 x 10^4^ cells/well in a 24-well plate (Falcon). Following overnight adherence, the media in all test wells was changed to fresh media. The cells were then pre-treated for 2h with either a DPBS control, 500 ng/mL of 700 bp GFP dsRNA, or 500 ng/mL of 700 bp mCherry dsRNA at 37°C with 5% CO_2_. Pre-treatment for 2h was selected after completing a time course experiment to determine the optimal amount of time to pre-treat cells with dsRNA to induce viral knockdown (supplementary figure S1). All test wells were then exposed to VSV-GFP at a multiplicity of infection (MOI) of 0.1 and allowed to incubate for 24h at 37°C with 5% CO2 before supernatants were collected for TCID50 quantification as described above.

### 4.7 Elucidating the role of sequence length in dsRNAi

To explore the impact that the dsRNA sequence length had on the observed viral knockdown, dsRNA was synthesized to GFP that ranged in size from 200 bp to 700 bp and tested for ability to induce knockdown. THF or SNB75 were seeded in a 24-well plate at a density of 5.0 x 10^4^ cells/well. Following overnight adherence followed by a media change, the cells were pre-treated for 2h with either a DPBS control, 500 ng/mL of mCherry dsRNA, or 500 ng/mL of GFP dsRNA at lengths of 200 bp, 300 bp, 400 bp, 500 bp, 600 bp and 700 bp at 37°C with 5% CO_2_. All test wells were then exposed to VSV-GFP at an MOI of 0.1 and allowed to incubate for 24h at 37°C with 5% CO2 before supernatants were collected for TCID50 quantification as described above.

### 4.8 Use of viral genes for dsRNAi

#### 4.8.1 VSV-GFP

DsRNA was synthesized to the VSV viral genes of N protein and M protein as described above. Either THF or SNB75 were seeded in a 24-well plate at a density of 5.0 x 10^4^ cells/well. Following overnight adherence, the media in all test wells was changed to fresh media. The cells were then pre-treated for 2h with either a DPBS control, 500 ng/mL of VSV N protein dsRNA, 500 ng/mL of VSV M protein dsRNA, 500 ng/mL of mCherry dsRNA or a combination of 250 ng/mL of VSV N protein and 250 ng/mL of VSV M protein (500 ng/mL total of dsRNA) at 37°C with 5% CO_2_. All test wells were then exposed to VSV-GFP at an MOI of 0.1 and allowed to incubate for 24h at 37°C with 5% CO_2_ before supernatants were collected for TCID50 quantification as described above.

#### 4.8.2 HCoV-229E

DsRNA was synthesized for HCoV-229E viral genes of RdRp, Spike protein, N protein and M protein as described above. MRC5 cells were seeded in a 24-well plate at a density of 7.5 x 10^4^ cells/well. Following overnight adherence, the media in all test wells was changed to fresh media. The cells were then pre-treated for 2h with either a DPBS control, 500 ng/mL of 229E RdRp, 500 ng/mL of 229E Spike protein, 500 ng/mL of 229E N protein dsRNA, 500 ng/mL of 229E M protein dsRNA or 500 ng/mL of mCherry dsRNA at 37°C with 5% CO_2_. All test wells were then exposed to HCoV-229E at an MOI of 0.02 and allowed to incubate for 24h at 37°C with 5% CO_2_ before supernatants were collected for TCID50 quantification as described above.

#### 4.8.3 SARS-CoV-2

DsRNA was synthesized for the SARS-CoV-2 viral genes, N protein and M protein, as described above. Calu-3 cells were seeded in a 12-well plate at a density of 2.0 x 10^5^ cells/well. Two days later, the media was replaced with fresh media. The cells were then pretreated for 2h with either 1000 ng/mL of mCherry dsRNA control, 1000 ng/mL of SARS-CoV-2 M protein dsRNA or 1000 ng/mL of SARS-CoV-2 N protein dsRNA at 37°C with 5% 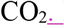Following pre-treatment, the cells were exposed to SARS-CoV-2 at an MOI of 1.0 for 1h, washed twice with sterile 1x PBS, and the dsRNA added back to the appropriate wells. After 24h, total RNA isolation was performed using the RNeasy Mini Kit (Qiagen) according to the manufacturer’s protocol. SARS-CoV-2 specific genome levels were measured by qPCR using SsoFast EvaGreen supermix (Bio-Rad) according to manufacturer’s protocol.

### 4.9 dsRNAi in human primary Bronchial/Tracheal Epithelial Cells (pBECs)

#### 4.9.1 Culture of human pBECS

Normal human primary bronchial/tracheal epithelial cells (pBECs) were purchased from ATCC (PCS-300-010). The pBECs were transferred to six T25 flasks containing complete Airway Epithelial Cell medium (ATCC) and cells were incubated at 37°C with 5% CO_2_ until they reached approximately 80% confluence. Cells were then detached using 0.25% trypsin (Gibco) and transferred to 1 μm transwell permeable supports (Falcon) in a 24-well plate at a density of 3.3 x 10^4^ cells/insert (200 μL per insert). The basolateral side received 700 μL of complete Airway Epithelial Cell medium. Media changes were made every 2d with the apical layer receiving 200 μL and the basolateral layer receiving 700 μL. Once the cells reached 100% confluence, the media was aspirated from the transwell which was then transported to a new 24-well plate and 600 μL of PneumaCult™ALI Maintenance medium (STEMCELL Technologies) was added to the basolateral side. Cells were left to grow for 28d with basolateral media changes occurring every 2d. After approximately 7d, apical washes using 200 μL of DPBS were performed every week to clear the cells of mucus production. After 28d the cells were used for experiments.

#### 4.9.2 Soaking with long dsRNA

Transwells containing the 28-day cultured pBECs were washed once by incubation with with 200 μL of sterile DPBS for 40 min. The transwells were then moved to a new 24-well plate wherein each test well contained 600 μL of fresh PneumaCult™-ALI Complete Base Medium (STEMCELL Technologies) for the basolateral side. The DPBS was removed from the apical side of the test transwells and were then exposed to either media alone, 500 ng/mL of dsRNA (VSV N protein, HCoV-229E M protein or mCherry as a control) or 50 μg/mL of pIC for 2h at 37°C with 5% CO_2_. Following the 2h incubation, appropriate test wells were exposed to either VSV-GFP (MOI = 0.1) or HCoV-229E (MOI = 0.1) and incubated for 24h before the supernatants were collected and the TCID50 was quantified as described above.

### 4.10 Soaking versus transfection with siRNA

In order to directly compare the effects of soaking with long dsRNA or siRNA on virus inhibition, THF and SNB75 cells were seeded in 24-well plates at a density of 5.0 x 10^4^ cells/well. Following overnight adherence and a fresh media change, cells were exposed to either a DPBS control, 2 nM of 700 bp GFP dsRNA, 2 nM of GFP Silencer^®^ siRNA (Ambion) or 2 nM of the negative control Silencer^®^ siRNA (Ambion) for 2h at 37°C with 5% CO_2_. Cells were exposed to nanomolar concentrations (equivalent to 500 ng/mL of 700 bp dsRNA) to ensure that the same number of dsRNA and siRNA molecules were added in each treatment group. Following this incubation, wells were exposed to VSV-GFP at an MOI of 0.1 and incubated for 24h at 37°C with 5% CO2 before supernatants were collected for TCID50 quantification as described above.

For validation that the siRNA molecules were functional and capable of inducing knockdown, the siRNA molecules were transfected into SNB75 and THF cells and subsequent viral numbers were quantified. THF and SNB75 cells were seeded 24-well plates at a density of 5.0 x 10^4^ cells/well. Following overnight adherence and a fresh media change, cells were 10 nM of GFP Silencer^®^ siRNA (Ambion) or 10 nM of the negative control Silencer^®^ siRNA (Ambion) was transfected into the cells using Lipofectamine RNAiMAX (Invitrogen). Cells were transfected with 10nM siRNA as recommended by the manufacturer. Following a 24h incubation at 37°C with 5% CO_2_, wells were washed twice with DPBS and then exposed to VSV-GFP at an MOI of 0.1 and incubated for 24h at 37°C with 5% CO2 before supernatants were collected and the TCID50 was quantified as described above.

### 4.11 Inducing viral inhibition using combination dsRNA molecules that target multiple viral genes

To determine whether combination dsRNA could induce viral knockdown via inhibition of multiple viral genes at once, THF cells were seeded at a density of 5.0 x 10^4^ cells/well in a 24-well plate. Following overnight adherence, the media in all test wells was changed to fresh media. The cells were then pre-treated for 2h with either a DPBS control, 500 ng/mL of 5’N-3’M (first 350 bp are VSV N protein and last 350 bp are VSV M protein), 500 ng/mL of 5’M-3’N (first 350 bp are VSV M protein and last 350 bp are VSV N protein), 500 ng/mL of N-M Alt (50 bp of VSV N protein and 50 bp of VSV M protein in alternating fashion for 700 bp), or 500 ng/mL mCherry dsRNA at 37°C with 5% CO2. All test wells were then exposed to VSV-GFP at an MOI of 0.1 and allowed to incubate for 24h at 37°C with 5% CO_2_ before supernatants were collected for TCID50 quantification as described above. The cell monolayers were collected in Trizol so that total RNA could be extracted and cDNA was synthesized as described above in *section 2.4*. The expression of VSV genes (M and N protein) was measured by qRT-PCR using the same method as outlined above in *section 2.4*. The sequences and accession number for the primer sets used here are outlined in **Table 2**. The VSV gene expression of cells exposed to the 5’M-3’N molecule was not measured due to the small size of the M protein gene which made it impossible to develop qPCR primers that did not amplify a region of the M-N dsRNA that was used to soak the cells.

### 4.12 Dicer knockout studies

For successful knockdown, the RNAi pathway requires the use of Dicer to cleave viral RNAs into siRNAs. To provide evidence that the knockdown observed here was due to RNAi, a Dicer1 knockout mouse MSC cell line (Dicer1 -/-) was used along with its corresponding functional Dicer1 cell line (Dicer1 f/f). For each experiment, both the knockout and functional Dicer MSC cell lines were seeded at a density of 5.0 x 10^4^ cells/well in a 24-well plate. Following overnight adherence, the media in all test wells was changed to fresh media. Both cell types were then pre-soaked for 2h with either a DPBS control, 500 ng/mL of VSV N protein dsRNA, or 500 ng/mL of mCherry dsRNA at 37°C with 5% CO_2_. All test wells were then exposed to VSV-GFP at an MOI of 0.1 and allowed to incubate for 24h at 37°C with 5% CO_2_ before supernatants were collected for TCID50 quantification as described above.

### 4.13 Statistical analyses

All data sets were tested for a normal distribution (Shapiro-Wilk) and homogeneity of variance (Levene’s) using R and RStudio (R Core Team, 2014; RStudio Team, 2015). Further statistical analyses were also completed using R and RStudio. For the viability, VSV gene expression and viral titer data, a one-way analysis of variance (ANOVA) was completed followed by a Tukey’s post-hoc test to compare between all exposure conditions. When determining whether the IFN genes were upregulated, a one-way ANOVA was completed followed by a Dunnett’s multiple comparisons post-hoc test to detect significant differences from the control condition. With the siRNA transfection data, a two-tailed unpaired t-test was completed. For all statistical analyses, a p-value less than 0.05 was considered significant. All data is presented as the average of experimental replicates + SEM.

## Acknowledgements

The authors would like to acknowledge Dr. Tamiru Alkie for his help with establishing the primary bronchial epithelial/tracheal cell cultures.

## Author Contributions

RJ performed the experiments involving SARS-CoV-2 and the associated analyses. KM contributed to experimental design and funding of the SARS-CoV-2 work. SS performed the remaining experiments, all of the associated analyses, contributed to experimental design and wrote the first draft of the manuscript. SDO contributed to experimental design, funding of the project, and writing of the manuscript. All authors contributed to manuscript revisions and approved the final submitted version.

---

**Supplementary Figure S1.**
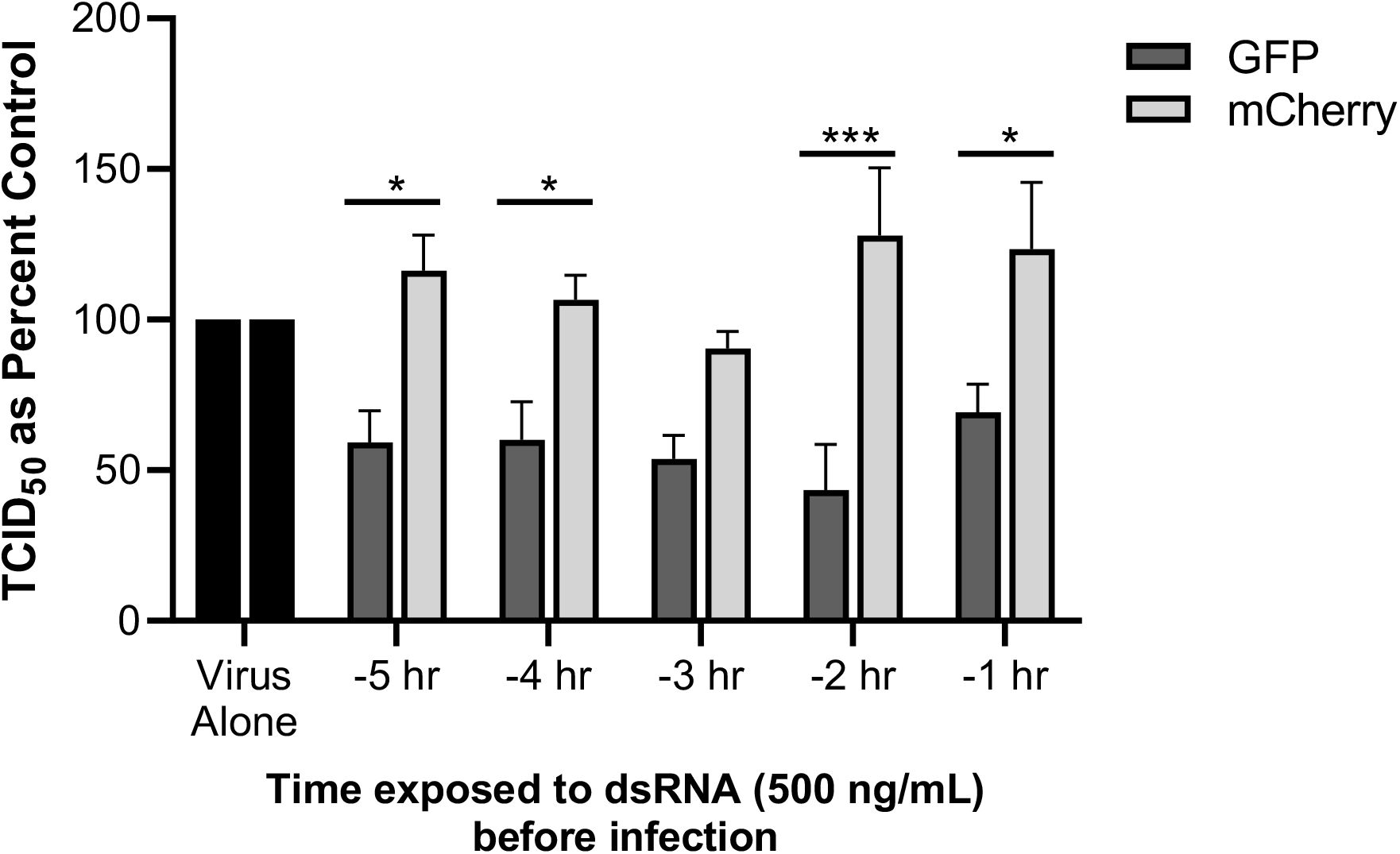
dsRNA effective at limiting virus VSV-GFP replication 1-5h prior to infection and at the time of infection. M14 Cells (75,000 cells/well) were exposed to 500 ng/mL of each dsRNA (700 bp each) at various times before infection with VSV-GFP (MOI = 1). Following 24 hours of infection, supernatants were collected and the TCID50 was calculated using HEL-299 cells. This has been repeated three times. Significant differences were assessed between mCherry and GFP at each individual timepoint using a Sidak’s multiple comparisons test.

